# Unveiling the multi-step solubilization mechanism of single vesicles by detergents

**DOI:** 10.1101/638189

**Authors:** Paul A. Dalgarno, José Juan-Colás, Gordon J. Hedley, Lucas Piñeiro, Mercedes Novo, Cibran Perez Gonzalez, Ifor D. W. Samuel, Mark C. Leake, Steven Johnson, Wajih Al-Soufi, J. Carlos Penedo, Steven D. Quinn

## Abstract

The solubilization of membranes by detergents is critical for many technological applications and has become widely used in biochemistry research to induce cell rupture, extract cell constituents, and to purify, reconstitute and crystallize membrane proteins. The thermodynamic details of solubilization have been extensively investigated, but the kinetic aspects remain poorly understood. Here we used a combination of single-vesicle Förster resonance energy transfer (svFRET), fluorescence correlation spectroscopy and quartz-crystal microbalance with dissipation monitoring to access the real-time kinetics and elementary solubilization steps of sub-micron sized vesicles, which are inaccessible by conventional diffraction-limited optical methods. Real-time injection of a non-ionic detergent, Triton X, induced biphasic solubilization kinetics of surface-immobilized vesicles labelled with the Dil/DiD FRET pair. The nanoscale sensitivity accessible by svFRET allowed us to unambiguously assign each kinetic step to distortions of the vesicle structure comprising an initial fast vesicle-swelling event followed by slow lipid loss and micellization. We expect the svFRET platform to be applicable beyond the sub-micron sizes studied here and become a unique tool to unravel the complex kinetics of detergent-lipid interactions.

## Introduction

Detergent-induced membrane solubilization is critical for applications including membrane-protein purification^1, 2^, and targeted drug delivery, where vesicle rupture enables release of encapsulated therapeutics^3^. Despite decades of widespread use, the complexity of membrane solubilization, coupled with limitations in current technology, have made characterizing its mechanism extremely challenging.^4^

Initial biochemical experiments indicated that the rate of membrane solubilization depends on the lipid phase, and the type and concentration of detergent^5^. The non-ionic detergent Triton X 100 (TX-100), for example, solubilizes phosphocholine (PC) rich membranes relatively slowly below the gel-to-liquid transition temperature but speeds up rapidly in the fluid phase^6, 7^. For most biochemical applications, TX-100 is the solubilizer of choice, and is used as a reference for measuring the activity of other surfactants^8, 9^. Turbidity measurements also reported the TX-100:lipid ratios required to solubilize lipid vesicles as a function of phase^10, 11, 12^ and lipid^13^, and isothermal titration calorimetry has probed the initial TX-100-membrane interaction^14^. These experiments suggest an interplay between surfactant monomers and lipids at the detergent’s critical micellar concentration in which lipid re-arrangement leads to heat transfer and mixed-micelle formation within the intact membrane^15^.

Importantly, the solubilization activity of TX-100 is inhibited by membrane cholesterol, though the precise reason for this is unknown. Cholesterol may initiate liquid ordered, detergent-resistant regions across the membrane^16, 17^, but whether these are microdomains^18^, detergent-resistant rafts^19^ or a combination of both^20^ requires confirmation. Alternatively, TX-100 may promote liquid-ordered phases via interaction with order-preferring cholesterol-rich regions rather than initiating lipid reorganization^21^.

Despite its complexity, the most widely adopted model to describe membrane solubilization is the three state mechanism^22^. In State 1, detergent monomers partition the bilayer until saturation is reached^23, 24^. State 2 involves the formation of mixed detergent-lipid micelles coexisting with the bilayer, and State 3 corresponds to breakdown of the membrane into mixed-micelles in solution. There are, however, many unanswered questions, particularly regarding the timescales of these processes and whether they occur sequentially or are interconnected, and ambiguities left on the mechanism are due to a lack of methods that can capture the entire process. While Cryo-TEM, NMR and conventional dynamic light scattering all provide snapshots of the membrane conformation^25, 26, 27^, they cannot provide dynamic insight. Conversely, ITC probes the thermodynamics and turbidity measurements reveal solubilization conditions, but neither reveal structure^28, 29^. Molecular dynamics simulations have attempted to bridge this gap^17, 30, 31^ and coarse-grained simulations reveal a sequence of events in broad agreement with the three-state mechanism^32^. Additionally, phase contrast and fluorescence microscopy have also been used to study the solubilization mechanism of giant (10-20μm) unilamellar vesicles^33^. However, conventional optical microscopy is diffraction limited and therefore cannot monitor solubilization processes for vesicles smaller than ∼250 nm, which are most commonly used for biotechnological applications. Furthermore, traditional optical microscopy quantifies macroscopic changes in size and packing density but provides little structural information at the molecular level. Therefore, there is a need to develop structural imaging methods that, while preserving the ability to monitor individual vesicles interacting with detergents in real time, can probe inter-molecular interactions.

Here we have explored the use of single-vesicle Förster resonance energy transfer (svFRET) as an imaging modality to study rapid structural changes in sub-micron vesicles in response to TX-100. FRET is sensitive to 1-10 nm distances between two small organic dyes termed donor and acceptor.^34^ In svFRET, these dyes are incorporated directly within the membrane and act as a macro-reporter of molecular interaction. Although svFRET has been applied to investigate the kinetics of membrane fusion^35, 36^ and pore formation^37^, its potential for characterizing solubilization kinetics has not been reported.

## Results and Discussion

PC- and phosphoserine (PS)-rich model-membrane vesicles incorporating 20% cholesterol were prepared as detailed in the **Supplementary Methods** and are schematically shown in **Figure S1**. The amounts of donor (Dil) and acceptor (DiD) per vesicle were optimized (1:1, 0.1 % of each dye) such that the average FRET efficiency (E_FRET_) per vesicle was initially close to 0.5, enabling nanometer length scale changes to be quantified by an observable change in E_FRET_ in either direction. The production of homogeneously distributed unilamellar vesicles was confirmed by dynamic light scattering (**Figure S2**). Steady-state fluorescence measurements were carried out as an initial step to characterize the interaction between TX-100 and labelled vesicles. As the concentration of TX-100 was progressively increased, we observed a decrease in E_FRET_ (**Figure 1a**), from a value of 0.43 ± 0.05 in the absence of TX-100, to 0.13 ± 0.02 in the presence of 4.4 mM with a half-maximal concentration constant of 0.39 ± 0.07 mM (see **Supplementary Methods** for FRET calculation details). These data suggest that the addition of TX-100 induces changes in vesicle structure, or composition, that results in a high distance separation between the dyes. The decrease in E_FRET_ was further confirmed by time-correlated single photon counting, where the amplitude weighted average lifetime of Dil progressively increased as a function of TX-100 (**Figure 1b, Table S1**).

**Figure 1.**
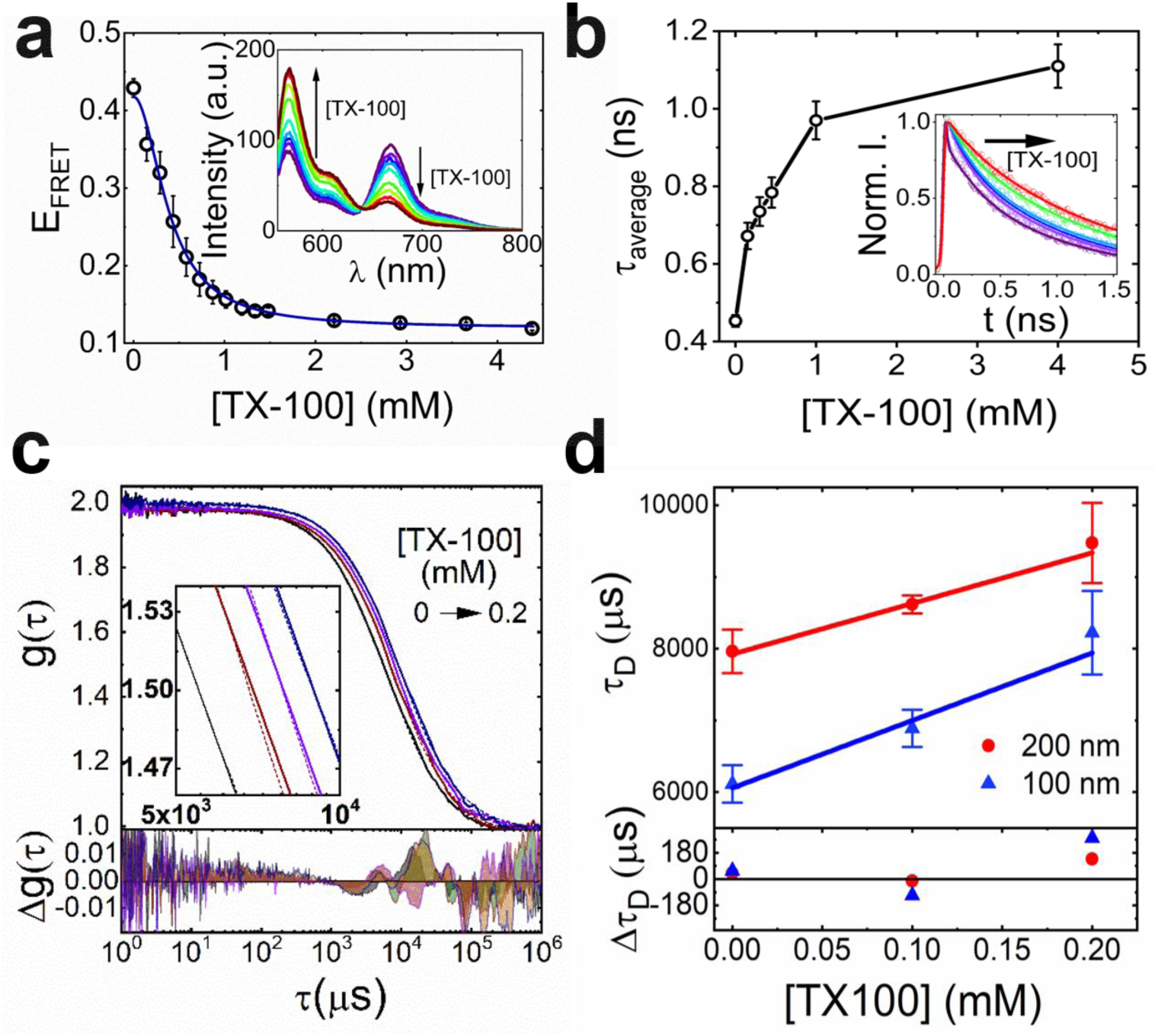
TX-100 vesicle interactions reported using ensemble FRET and FCS. (a) FRET efficiency of Dil/DiD labeled vesicles versus TX-100. The solid line represents a Hill model fit. Inset: corresponding variation in fluorescence spectra. (b) Average lifetime of Dil versus TX-100. Inset: corresponding time-resolved fluorescence decays. Solid lines represent bi-exponential fits. (c) Top: FCS cross-correlation curves (solid lines), fits (dashed lines) (inset: zoomed in) associated with 100 nm-(black) and 200 nm-sized (dark red) vesicles. Also shown are FCS curves for 200 nm-sized vesicles in the presence of 0.1 mM (purple) and 0.2 mM (blue) TX-100. Bottom: residuals of the fits. (d) Top: Diffusion times of NBD-PC labeled vesicles as a function of TX-100. Solid lines represent linear fits. Bottom: corresponding residuals.

Having established FRET as a sensor of fluorophore separation in the ensemble, fluorescence correlation spectroscopy (FCS) was used to probe the diffusion of single vesicles. The high sensitivity of FCS to the size of diffusing vesicles makes it an attractive technique for accessing their diameter under solubilizing conditions^38, 39, 40^. The translational diffusion times of vesicles prepared with 0.2 % (1-palmitoyl-2-{6-[(7-nitro-2-1, 3-benzoxadiazol-4-yl) amino] hexanoyl}-*sn*-glycero-3-phosphocholine) (NBD-PC) (**Figure S3)**, a fluorescent analog of PC, were recorded. NBD-PC replaced Dil and DiD for compatibility with the apparatus. Normalized cross-correlation functions obtained from freely-diffusing vesicles progressively shifted towards longer diffusion times as a function of TX-100 (**Figure 1c**). In the absence of detergent, vesicles of diameters ∼100 nm and ∼200 nm displayed diffusion times, τ_D_, across the confocal volume of 6.2 ± 0.2 ms and 7.8 ± 0.2 ms, respectively (**Figure 1d**). τ_D_ associated with the diffusion of 100 nm-diameter vesicles increased by 12 % in 0.1 mM TX-100 and by 34 % in 0.2 mM TX-100. The larger 200 nm-diameter vesicles also displayed a similar trend, representing an 8 % and 22 % increase in hydrodynamic diameter at 0.1 mM and 0.2 mM TX-100, respectively. These data point toward an increase in mean vesicle diameter when incubated with TX-100, and was attributed to vesicle expansion, fusion or a combination of both in solution.

To rule out the possibility of fusion and investigate in more detail each step of the solubilization process, Dil/DiD labelled vesicles containing a low percentage of biotinylated lipids and 20% cholesterol were immobilized onto a NeutrAvidin-coated surface imaged via total internal reflection fluorescence microscopy. As illustrated in **Figure 2a**, biotinylated vesicles were anchored to NeutrAvadin tethered to the surface via biotinylated polyethylene glycol (PEG). In the absence of TX-100 the vesicles were stable with no variation in the svFRET efficiency observed (**Figure 2b**). Perturbation of single vesicles by TX-100 was then reported as observable changes in the svFRET efficiency in real-time with 50 ms time integration (**Figure 2b, 2c**). To suppress photo-bleaching and optimize conditions for svFRET, the fluorescence response of single vesicles labelled with DiD were investigated as a function of excitation intensity and percentage of dye-loading content. As demonstrated in **Supplementary Text I, Figure S4** and **Table S2**, excitation intensities < 0.04 mW/cm^2^ with 0.25 % dye were necessary for long-term (180 s) stability of the incorporated dyes. In the absence of TX-100, the FRET efficiency from single vesicles remained largely invariant with a value of E_FRET_ ∼ 0.5. Injection of 0.16 mM TX-100 then induced variations in E_FRET_ and total intensity on remarkably different time scales. Immediately after injection of TX-100, the FRET efficiency decreased from a value of ∼0.46 to ∼0.22 in a 250 sec time window (**Figures 2d, S5**). Most of this change happened in the first 100 seconds after injection, pointing towards a ∼20% increase in the separation distance between FRET pairs. Assuming spherical vesicles, this distance scales directly with the vesicle radius and thus agrees well with the FCS data. Importantly, within this time window, the total intensity decreased only by 7%. At time scales longer than 250 seconds, the FRET efficiency did not change further, whereas the total intensity decreased progressively to ∼ 50% of its initial value. The different timescales and responses of both signals suggest that they represent different distortions of the vesicle structure. The rapid decrease in FRET efficiency without significant variation in total intensity indicates a structural change involving no loss of lipid content and we assigned it as arising from vesicle swelling induced by TX-100 molecules inserting into the lipid bilayer and increasing the average inter-dye distance. The time window where the FRET efficiency does not change but the total intensity is strongly decreased suggests a structural distortion involving the diffusion of lipids into solution and it was assigned to a lysis step resulting in the formation of micelles. The FRET efficiency plateau value observed in this time window (E ∼ 0.22) represents an inter-dye distance ∼6.5 nm within these micelles. These observations point towards a fast vesicle expansion event with half-life t^E^, followed by a slower lysis event (t^L^). The expansion step was observed to occur on average 12 times faster than lysis, and at 0.08 mM TX-100, the half-lives associated with each event increased by ∼75 % (**Figure 2e**).

**Figure 2.**
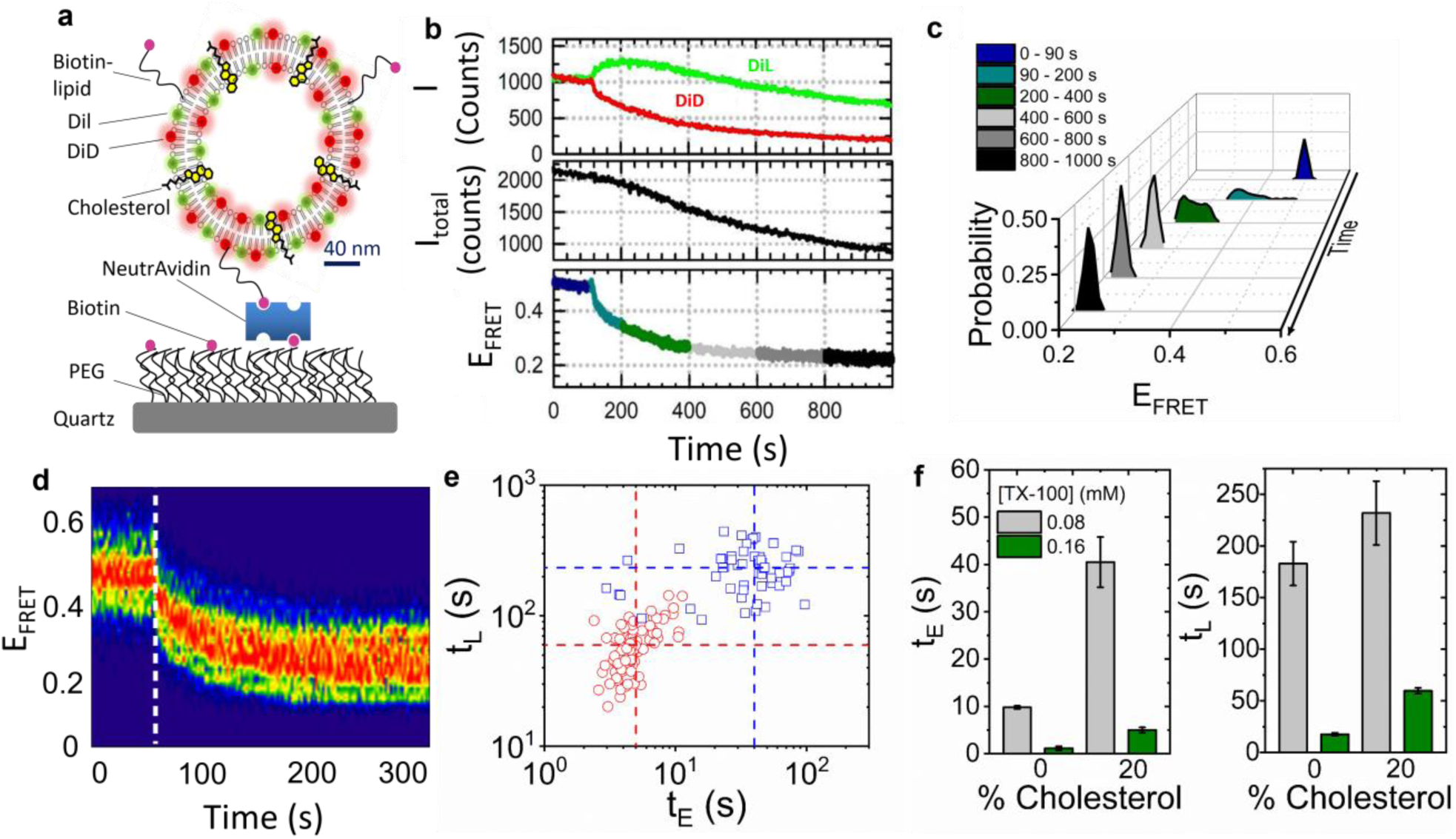
Real-time visualization of solubilization kinetics by svFRET. (a) Schematic of the immobilization scheme. (b) Representative variation in the fluorescence emission of Dil and DiD (top panel), the sum of their intensities (middle panel) and the corresponding variation in FRET efficiency obtained before (< 90 s) and after (>90 s) injection of 0.16 mM TX-100. (c) Relative FRET state occupancies observed over 1000 s. (d) FRET contour plot showing the variation in E_FRET_ before and after TX-100 injection (dashed white line) (N = 105). (e) Corresponding scatter plot of expansion half-live, t_E_ versus that of lysis, t_L_ obtained after injection of 0.08 mM (blue) and 0.16 mM (red) TX-100. Dashed lines represent the center of each distribution. (f) Comparative bar plots summarizing the variation in t_E_ and t_L_ as a function of TX-100 and percentage of cholesterol incorporated within the vesicle bilayer. Error bars indicate the standard error of the mean.

Since cholesterol alters the membrane structure, we next employed svFRET to assess the influence of cholesterol on the stability and kinetic mechanism of vesicle solubilization. 200 nm diameter vesicles were prepared in the absence of cholesterol and were induced to solubilize by 0.08 mM TX-100 (**Figure S6**). Here, the expansion and lysis half-lives reduced by ∼88 % and ∼21 % respectively, compared with vesicles loaded with 20 % cholesterol. When the TX-100 concentration was doubled, t_E_ and t_L_ reduced further to 1.2 ± 0.4 and 17.6 ± 1.6 seconds, respectively, as shown via a comparative bar plot summarizing the relative variations in t_E_ and t_L_ as a function of cholesterol and TX-100 concentration (**Figure 2f**).

Although TX-100 induced vesicle expansion is clear from the svFRET assay, quantifying the extent of TX-100 incorporation and mass loss is more complex. To assess this, a complementary label-free quartz-crystal microbalance with dissipation (QCM-D) monitoring approach was employed (**Supplementary Text II**). Here, PC/PS vesicles containing 20 % cholesterol were immobilized onto a quartz sensor surface and interactions with 0.16 mM TX-100 were followed by changes in oscillation frequency and dissipation, reflecting mass and viscoelasticity on the sensor surface, respectively (**Figure 3a**). Interactions between immobilized vesicles and TX-100 were observed via changes in both frequency and dissipation traces immediately after TX-100 injection corresponding to a ∼5% mass gain at the sensor which we attributed to TX-100 incorporation into vesicles. This was followed by a non-destructive interaction that leads to a conformational change in intact vesicles over the first 35 s. As the local TX-100 concentration then increased, the deposited mass accumulated on the surface leading to a decrease in resonance frequency. A substantial mass loss of ∼63% over the following 85 s was then observed via an increase in resonance frequency, indicating material immobilized to the surface was released into solution (**Figure 3b**). Control experiments performed simultaneously indicated no interaction with the PEG-coated sensor surface and TX-100 (**Figure 3a**).

**Figure 3.**
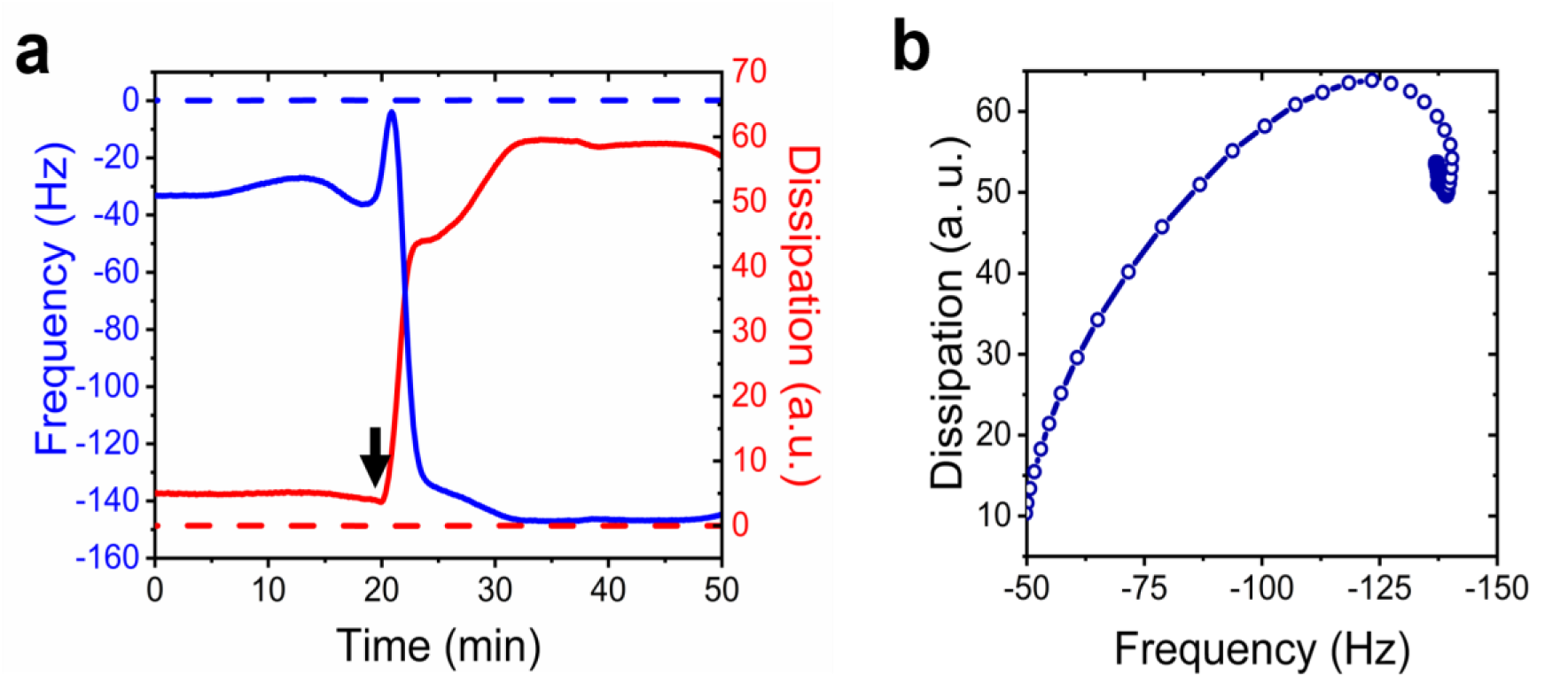
TX-100 induced vesicle solubilization monitored by QCM-D. (a) Variation in frequency (blue) and dissipation (red) of the 7^th^ overtone associated with surface immobilized vesicles in the presence of TX-100. The dashed lines represent data collected from a control sensor pre-treated with PEG and NeutrAvidin, but lacks vesicles. The arrow indicates the time-point of the solubilization. (b) Frequency versus dissipation observed during the interaction between surface immobilized vesicles and TX-100.

These findings support a mechanism through which TX-100 accumulates on the curved membrane surface, preceding a rapid 20 % expansion of the vesicle structure that, in turn, precedes a slower lysis event (**Figure 4**). It is also possible that the curvophilic nature of TX-100 may, in addition, promote permeabilization, contributing to content leakage^12, 32, 41^.

**Figure 4.**
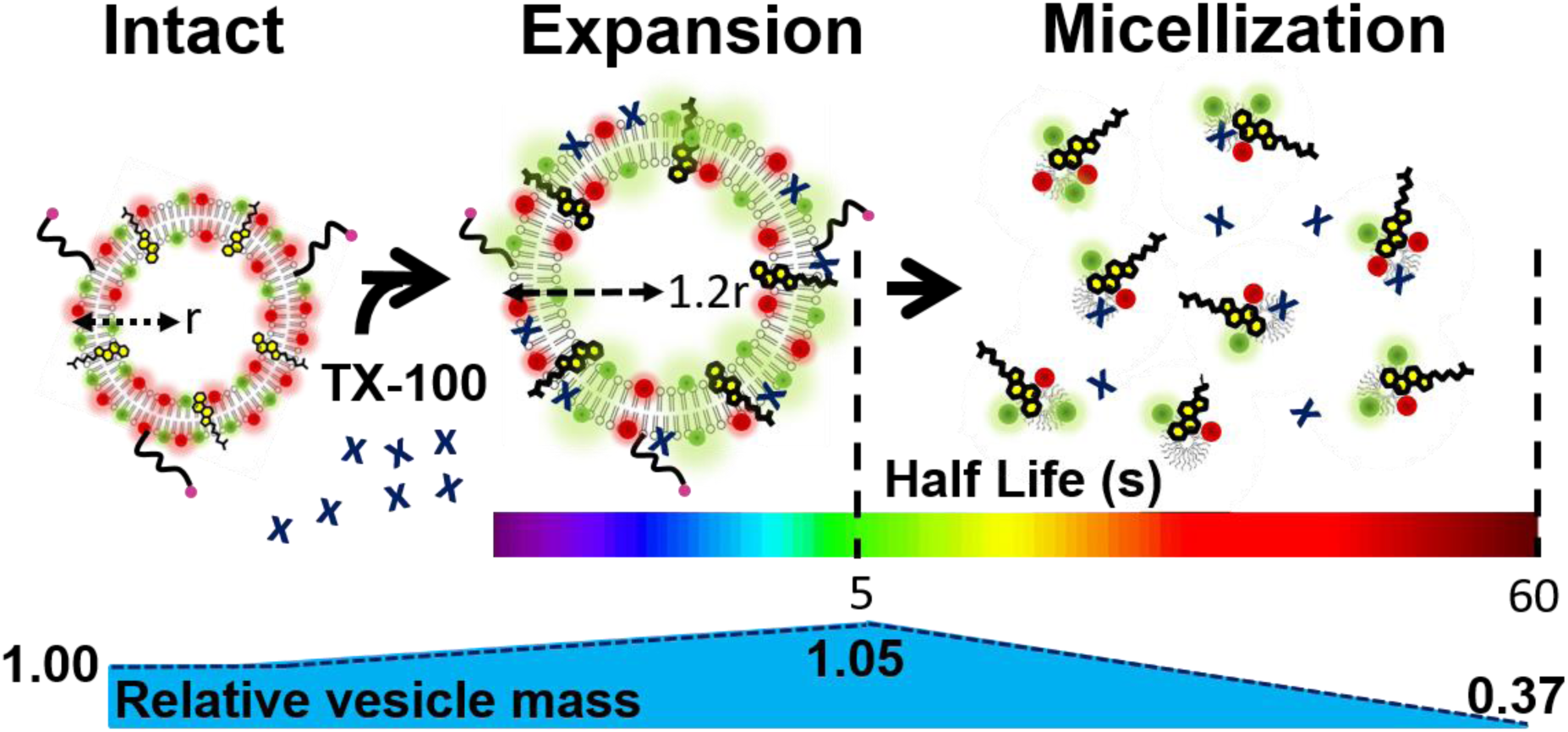
Mechanism of TX-100 induced vesicle solubilization. Detergent molecules approach lipid vesicles inducing a fast conformational expansion prior to lysis and the release of mixed-micelles into solution.

We have directly observed the solubilization of model-membrane vesicles in response to TX-100 using svFRET, FCS and QCM-D. Taken together, these trajectories separate each stage of the solubilization pathway, enabling kinetic parameters to be assigned to each process. We report TX-100 induced solubilization as a sequence of events, whereby TX-100 deposition onto the vesicles induces vesicle expansion prior to micellization. The large changes in solubilization kinetics for a vesicle model as a function of cholesterol suggest that the membrane constituents are important regulators of membrane stability. Although the experimental data discussed here were obtained with TX-100, the conclusions should be applicable to a wide variety of detergents and may lead to more effective and efficient applications involving membrane protein purification.

## ASSOCIATED CONTENT

Figures S1-S6, Tables S1-S2, methods and text are found in the Supporting Information.

## Acknowledgements

We thank Dr. Arvydas Ruseckas for helping with fluorescence lifetime measurements and the EPSRC (EP/P030017) for support.

