## Supplementary Information for "Unveiling the multi-step solubilization mechanism of single vesicles by detergents"

### Methods

#### Materials

1-palmitoyl-2-oleoyl-sn-glycero-3-phospho-L-serine (sodium salt) (PS), 1-palmitoyl-2-oleoyl-glycero-3-phosphocholine (PC) and 1,2-dioleoyl-sn-glycero-3-phosphoethanolamine-N-(cap biotinyl) (sodium salt) (biotinylated lipid) phospholipids were purchased from Avanti Polar Lipids Inc. 1,1'-Dioctadecyl-3,3,3',3'-Tetramethylindocarbocyanine Perchlorate (DiI) and 1,1'-Dioctadecyl-3,3,3',3'-Tetramethylindodicarbocyanine, 4-Chlorobenzenesulfonate Salt) (DiD) membrane stains were purchased from ThermoFisher Scientific.1-palmitoyl-2-{6-[(7-nitro-2-1,3-benzoxadiazol-4-yl)amino]hexanoyl}-sn glycerophosphocholine (NBD-PC) was purchased from Sigma Aldrich. All phospholipid samples were used without additional purification and stored in chloroform at -20$^{o}$C prior to use. DiI, DiD and cholesterol stocks were stored at 4$^{o}$C in chloroform prior to use. Triton X-100 was purchased from Sigma Aldrich and freshly suspended in 50 mM Tris (pH 8) prior to use.

#### Preparation of small unilamellar vesicles

Mixtures of lipids and lipophilic dyes were homogeneously dispersed in chloroform, dried by nitrogen flow and stored under continuous vacuum pumping at room temperature for 5 hours. Phospholipid mixtures were subsequently re-suspended in buffer solution (50 mM Tris, pH 8) and mixed well by vortex. Small unilamellar vesicles were prepared by the extrusion method at room temperature, in which they were passed through a polycarbonate membrane filter of defined pore size. A molar ratio of 65:35 PC:PS was used for FRET and FCS studies. Vesicles were labelled with dyes (0-0.5 %), cholesterol (0-20 %) and biotin (1 %) at the molar percentages specified in the text. The mean size of the prepared vesicles in solution was evaluated by dynamic light scattering using a Zetasizer μV molecular size detector (Malvern Instruments Ltd., UK).

#### Cy5-ssDNA Labelling Reaction, Precipitation and Purification

C20-3'amino DNA was labelled with the DiD analogue Cy5 (Integrated DNA Technologies, IA, USA) for single-molecule analysis. Here, the dry pellet of DNA was dissolved in deionised water to a final concentration of 25 μg/μL. Cy5 was suspended in dimethyl sulfoxide (DMSO) to a final concentration of 22.5 mM, dried using a SpeedVac system for 4 hours and re-suspended in a solution composed of 75% (v/v) labelling buffer (0.1 M sodium tetraborate, pH 8.5, 7% (v/v) deionised water, 14% (v/v) DMSO and 4 % (v/v) C20-3'amino DNA). This was stored in darkness at room temperature under gentle agitation for 16 hours. Sodium acetate (0.3 M) and ice cold absolute ethanol (250 μL) were subsequently added to the labelling reaction, mixed gently by inversion and incubated at -20°C for a further 16 hours. After incubation, the solution was centrifuged at 13000 rpm for 1 hour and the supernatant removed. The remaining pellet was rinsed with cold ethanol, dried with nitrogen and dissolved in 50 mM Tris-HCl buffer (pH 7.5). Denaturing polyacrylamide gel electrophoresis was subsequently performed at 4°C to separate labelled and unlabelled DNA. Final Cy5-DNA constructs were stored at 4°C prior to use.

**Steady-state fluorescence spectroscopy**

Fluorescence emission spectra were acquired using a Varian Eclipse fluorescence spectrophotometer. Spectra from DiI and DiD were recorded using an excitation wavelength of 532 nm at magic angle. FRET efficiencies were approximated by the apparent FRET efficiency, $E_{FRET} = (I_{665}/[I_{665} + I_{565}])$, where $I_{665}$ and $I_{565}$ represent the fluorescence intensities of the acceptor at 665 nm, and donor at 565 nm, respectively. The FRET efficiency data shown in Figure 1a was fitted to a Hill model of the form, $E_{FRET} = A + B\frac{[TX-100]^n}{k^n+[TX-100]^n}$, where A and B are the measured FRET efficiencies at the start and end of the titration, k is the half-maximal concentration constant and n is the Hill coefficient. The parameters of the fit shown in **Figure 1a** are A = 0.43 ± 0.04, B = 0.12 ± 0.01 , k = 0.39 ± 0.07 and n = 2.0 ± 0.3 ($\chi^2$ = 0.99). Error bars represent the standard error of the mean from 3 individual experimental runs.

**Time-resolved fluorescence spectroscopy**

Fluorescence lifetime measurements were performed with a Hamamatsu C6860 Synchroscan streak camera. The 80 MHz, 100 fs (full width half maximum) 800 nm output of a Ti:Sapphire oscillator was frequency doubled with a beta barium borate crystal, giving 400 nm excitation pulses. The 400 nm light, with an average power of less than 1 mW, was subsequently focussed through the optical path length

(1cm) of the solution cuvette. Fluorescence from the sample was then collected and collimated with a lens before being focussed onto the entrance slit of a Chromex 250is imaging spectrograph. Excitation light was removed with a yellow schott glass filter that cuts all light below 420 nm. A spectral window of 585–607 nm corresponding to DiI fluorescence emission was selected with the spectrograph before the light was directed into the streak camera. Time-resolved fluorescence dynamics were then recorded enabling time constants of approximately 10 ps to be resolved with instrument response deconvolution.

**Fluorescence Correlation Spectroscopy**

Samples were deposited on glass-bottomed well plates (Whatman) and excited by the linearly polarised light of a 488 nm continuous wave laser (Becker & Hickl) which was spectrally cleaned (Semrock, US, FF01-482/18), redirected by a dichroic mirror (Semrock, US, DI01-R488) and focused into the sample by a 60 x water immersion microscope objective (Olympus, UPLSAPO60xW/1.2) mounted in an inverted microscope (Olympus, IX-71). The fluorescence was focused onto a $\phi$ = 50 μm pinhole (Thorlabs) before being split by a 50:50 nonpolarising beamsplitter cube (Thorlabs). Each beam was then focused onto an avalanche photodiode (MPD50CTC APD, $\varnothing$ = 50 μm, Micro Photon Devices). An emission filter (Semrock, 525/45) placed in front of the beamsplitter was used to discriminate fluorescence from scattered light. The detector signals were processed and stored by two time-correlated single photon counting (TCSPC) modules (Becker & Hickl, SPC 132). Typically 20 million photons were collected for each correlation curve with count rates between 5 and 20 kHz. All measurements were made at a stabilized temperature of 25.0 ± 0.5 °C. The excitation power as measured in the focus of the microscope objective by a power meter (Thorlabs) was 0.02 mW corresponding to a mean irradiance of 7.15 kW/cm$^2$ assuming a Gaussian intensity distribution along the optical axis. The focal area and the detection volume were calibrated with Rhodamine 123 in aqueous solutions at low irradiance using an estimated diffusion coefficient of 4.6 ± 0.4 × $10^{-10}$ m$^2$ s$^{-1}$, yielding a radial 1/e$^2$ radius of $\omega_{xy}$ = 0.26 μm and volume of focus of V = 0.53 μm$^3$. Correlation functions were calculated according to $G = \frac{\langle I(t)+I(t+\tau)\rangle}{\langle I(t)^2\rangle}$ where I(t) is the intensity at time t and fitted according to $G = b_0 + \frac{1}{N}\left(1+\frac{t}{\tau_D}\right)^{-1}\sqrt{\left(1+\frac{t}{\Omega^2\tau_D}\right)^{-1}}\left(1+A_T e^{-t/\tau_T}\right)$ where N is the number of molecules, t is the correlation time, $A_T$ is the amplitude of the triplet and $\tau_T$ is the triplet time. $\Omega$ defines the ratio between the axial and radial $1/e^2$ radii, $\omega_z$ and $\omega_{xy}$, respectively: $\Omega = \frac{\omega_z}{\omega_{xy}}$. All diffusion coefficients were corrected for temperature and viscosity effects and are reported for 25°C. Power series were performed in order to determine the photobleaching limits. A triplet-state contribution of 1 μs with the expected irradiance-dependent amplitude was observed in all cases. All measurements were repeated at least 20 times and curves distorted due to occasional transits of big aggregates were excluded. The surfactant was added to the diluted vesicle samples immediately before the FCS measurement. Error bars indicate the standard error of the mean.

**Single-vesicle TIRF Spectroscopy**

Fluorescence emission at the donor and acceptor wavelengths were acquired from single vesicles by using a prism-type total internal reflection fluorescence microscope equipped with green (532nm) and red (635nm) lasers (Crystalaser, USA). Microscope slides were successively treated with biotinylated poly-ethyleneglycol (PEG) and NeutrAvidin, before pM concentrations of fluorescently-labelled vesicles were added. Fluorescence trajectories were acquired with an integration time of 50 ms. The base buffer used for imaging was 50 mM Tris (pH 8), 6 % (w/v) glucose, 165 U/mL glucose oxidase, 2170 U/mL catalase and 2 mM trolox. Specified concentrations of TX-100 were included in imaging buffer prior to being injected into the sample. Spatially-separated fluorescence images of donor and acceptor emission were collected in custom built relay optics with a 550 nm long-pass filter and imaged in parallel using an EMCCD camera (iXON, Andor Technology). All measurements were performed at room temperature. SvFRET efficiency after background correction was approximated by the apparent FRET efficiency, $E_{FRET} = (I_A/[I_A + I_D]) \sim R_o^6/([R_o^6+R^6])$, where $I_A$ and $I_D$ are the fluorescence intensities of the acceptor and donor, respectively, $R_o$ is the Forster radius and R is the separation between the probes. Since the quantum yields of DiI and DiD are similar, $E_{FRET}$ closely matches the true efficiency of energy transfer. Half-lives were calculated by applying double exponential fits consisting of a rise ($I = Ae^{t/tE}$) and decay ($I = Be^{-t/tL}$) component to the donor trajectories. Data analysis was carried out using laboratory-written analysis routines developed in

MATLAB 7. Ensemble information from svFRET measurements was obtained by assembling single-vesicle FRET trajectories into population FRET contour plots.

**Quartz Crystal Microbalance with Dissipation (QCM-D) monitoring.**

Quartz crystal microbalance with dissipation monitoring (QCM-D) experiments were performed using a Q-sense E4 system (Biolin Scientific). SiO2-coated AT-cut quartz sensors (QSX 303, Biolin Scientific) were used, for which the fundamental frequency was 4.95 ± 0.05 MHz. The sensors were initially subjected to a 10 minute cleaning step by UV−ozone, prior to being sonicated in solutions of 2 % Hellmanex III and 2 x ultrapure Milli-Q water for 10 minutes. The sensors were then dried with $N_2$ and placed under UV−ozone for a further 30 minutes. Each sensor was then immersed in 100% ethanol for 30 minutes and dried with $N_2$ before installation in the flow modules. The QCM-D flow chambers were first flushed with ultrapure Milli-Q water for 1 hour, and then with 50 mM Tris buffer (pH 8) for 20-30 minutes before each measurement until a stable baseline was established (<0.5 Hz shift over 10 min). The flow rate was kept constant at 20 µL/min. The sensor surfaces were then functionalized with biotinylated polyethyleneglycol (PEG) (Iris Biotech) which acts as a biocompatible support for specific immobilization of Avidin through the Avidin-Biotin interaction. The sensor surfaces were then rinsed with 50 mM Tris buffer (pH 8.0) for 15 min to remove unadsorbed molecules. Thereafter, Avidin was immobilized on the sensor surfaces by incubating a 0.1 mg/mL Avidin solution in 50 mM Tris buffer (pH 8.0) for 20 min, following a rinse step with 50 mM Tris buffer (pH 8.0) for 20 min to wash unbound Avidin molecules. Subsequently, vesicles coated with 1 % biotinylated lipids were immobilized on the sensor surfaces by incubation with a 33 µg/mL vesicle solution for 70 min. Triton X-100 (TX-100) detergent solutions at specified concentrations were then introduced into the QCM-D flow chambers. Changes in mass ($\Delta m$) were related to changes in frequency ($\Delta f$) via the Sauerbrey equation $\Delta m = -(C \cdot \Delta f)/n$ where n is the overtone number and C is a constant related to the properties of the quartz (17.7 ng $Hz^{-1}cm^{-2}$).

### Supplementary Text I

#### Optimization of Vesicle Photostability for svFRET Experiments

To optimize conditions necessary for FRET based vesicle-detergent experiments, the photobleaching rates of DiD-labelled PC/PS vesicles were investigated. The goal was to determine the factors that modulate acceptor photostability, and, to identify methods to minimise their effects. A schematic illustration of the vesicles is shown in **Figure S5 a**. The vesicles were studied using TIRF spectroscopy with $\lambda_{ex}$ = 635 nm as a function of excitation intensity (0.04 mW/cm$^2$, 0.4 mW/cm$^2$ and 0.8 mW/cm$^2$) and molar percentage of labelling (0.05 % - 0.5 %). In the absence of TX-100, DiD-labelled vesicles displayed photobleaching rates that were faster than those associated with single dyes, which we attributed to dye crowding on the membrane surface. For example, Cy5 fluorophores covalently attached to immobilized single-stranded DNA molecules remained photoactive for a period of ~60 seconds under an excitation intensity of 0.4 mW/cm$^2$ before displaying single-step photobleaching (**Figure S5 b**). In contrast, the fluorescence intensity from DiD-labelled vesicles was found to display increasing levels of destabilization as the excitation power increased from 0.04 to 0.8 mW/cm$^2$, though the effect was lessened by reducing the dye-loading on each vesicle from 0.5 % to 0.05 %. When the labelling density was 0.5 % and the excitation intensity set to 0.4 mW/cm$^2$, the DiD fluorescence trajectories displayed bi-exponential behaviour with an average decay time of 28 ± 4 s. As the excitation intensity decreased to 0.04 mW/cm$^2$, the fluorescence decays were dominated by longer lived components and an average decay time of 152 ± 7 s was observed. With a labelling concentration of 0.05 %, the average decay time at 0.4 mW/cm$^2$ was 35 ± 3 s and increased to 439 ± 10 s when the excitation power decreased ten-fold. At 0.8 mW/cm$^2$, the fluorescence trajectories decayed bi-exponentially with average decay times of 13 ± 1 s and 14 ± 1 s when labelling concentrations of 0.5 and 0.05 % were tested. Representative photobleaching traces of DiD labelled vesicles as a function of excitation intensity and dye loading are shown in **Figure S5 c-e**. Pre-exponential factors and kinetics of the photobleaching trajectories under the same conditions are shown in **Table S2**.

When imaging of labelled vesicles containing 0.125 % DiI and 0.125 % DiD, as necessary for smFRET experiments, was performed, an excitation intensity of < 0.04 $mW/cm^2$ was thus required for long-term photostability. Under this condition, the fluorescence trajectories of both dyes, and therefore the corresponding efficiency of energy transfer, were found to be invariant over the measurement time window as demonstrated by the representative fluorescence trajectories shown in **Figure S5 f, g**. An analysis of N > 500 vesicles revealed good photostability over a 3 minute time window, as demonstrated by a lack of variation in the FRET contour plot shown in **Figure S5 h.** Given that the effect of photobleaching was minimised, this provided an opportunity to study vesicle perturbation by TX-100 using svFRET as the optical sensor.

### Supplementary Text II

#### Vesicle solubilization monitored by Quartz Crystal Microbalance with Dissipation Monitoring (QCM-D)

Quartz crystal microbalance with dissipation monitoring (QCM-D) was employed to investigate conformational changes of vesicles immobilized to a gold coated quartz sensor substrate. Here, vesicles containing a base ratio of 65: 35 PC: PS incorporating 20 % cholesterol and 1 % biotinylated lipids were immobilized onto the QCM-D sensor surface as described in the Methods section. Interactions between the vesicles and TX-100 were monitored through real-time changes in oscillation frequency and dissipation, reflecting the mass and rigidity of the immobilized vesicles, respectively, as TX-100 was flushed over the sensor surface. Upon exposure of the vesicles to TX-100 concentrations < 0.16 mM solution, a small decrease in frequency was consistently observed in all harmonics which we attribute to the interaction of the detergent molecules with the lipid layer of the vesicles. At higher TX-100 concentrations, a pronounced interaction of the detergent with the surface-immobilized vesicles was observed as demonstrated by both frequency and dissipation responses at the sensor. The ripple observed in both frequency and dissipation traces immediately after TX-100 was introduced is attributed to two events: first, the deposition of TX-100 molecules on the vesicle membrane, and second, a non-disruptive interaction of the detergent molecules with the vesicles that leads to re-structuring of the vesicle conformation (**Figure 3a**). Control experiments performed simultaneously indicate no interaction with the PEG-coated sensor surface and TX-100 at the concentrations tested, as indicated by the lack of variation in the frequency and dissipation response after TX-100 injection. As TX-100 builds up, the deposited mass continues to increase leading to a decrease in resonance frequency. Subsequently, following a steep transition which indicates re-structuring of the vesicles, an increase in resonance frequency (i.e. decrease in mass) is observed in the harmonics of the resonant wave, indicating material immobilized to the surface, namely lipid and aqueous solution encapsulated by the vesicles, is released into the solution following TX-100 accumulation. The energy dissipation response supports these deductions, as prior to the steep transition there is an increase in energy dissipation, which relates to the rigidity decrease obtained from the deposition of TX-100 molecules. After a sharp, short transition, the dissipation levels are decreased indicating an increase in rigidity due to the incorporation of TX-100 onto the vesicles. This change in vesicle conformation also leads to a cross in the different harmonic dissipation trends of the sensor as they have a different penetration depth and therefore changes in conformation affect the energy dissipation response in different manners. The interaction of the TX-100 detergent at this concentration with the surface-immobilized vesicles can be observed more clearly by plotting the energy dissipation response against the change in resonance frequency for a given overtone (here the 7$^{th}$ overtone) as shown in **Figure 3b**. An interaction between TX-100 and immobilized vesicles can be deduced from the QCM-D sensing strategy whereby TX-100 binds to the surface-immobilized vesicles initiating a structural reorganization consistent with vesicle expansion, which precedes a major lysis event. When complete solubilization was achieved, the frequency and dissipation responses closely matched those obtained from a control sensor lacking vesicles.

**Table S1** Pre-exponential factors, fluorescence lifetimes and fitting errors associated with DiI in the presence of the acceptor DiD obtained from freshly prepared PC/PS vesicles containing 20 % cholesterol as a function of TX-100. Experimental conditions: Tris-HCl buffer, pH 7.9, 21°C.

| | **0 mM** | **0.15 mM** | **0.30 mM** | **0.45 mM** | **1.00 mM** | **4.00 mM** |
|---|---|---|---|---|---|---|
| **$a_1$** | 0.35 ± 0.02 | 0.29 ± 0.01 | 0.26 ± 0.01 | 0.27 ± 0.02 | 0.18 ± 0.01 | 0.12 ± 0.01 |
| **$\tau_1$ / ps** | 18 ± 1 | 140 ± 7 | 150 ± 7.5 | 200 ± 10 | 150 ± 7.5 | 121 ± 6 |
| **$a_2$** | 0.80 ± 0.02 | 0.71 ± 0.04 | 0.74 ± 0.04 | 0.73 ± 0.04 | 0.82 ± 0.04 | 0.88 ± 0.04 |
| **$\tau_2$ / ps** | 615 ± 7 | 890 ± 44.5 | 940 ± 47 | 1000 ± 50 | 1150 ± 58 | 1250 ± 63 |
| **$\tau_{av}$*/ ps** | ***453 ± 5*** | ***672 ± 34*** | ***735 ± 37*** | ***784 ± 39*** | ***970 ± 49*** | ***1110 ± 56*** |

* Amplitude weighted average lifetimes, $\tau_{av}$, were calculated according to the equation $\tau_{av} = \sum_{i=1}^{N} a_i \tau_i^2 / \sum_{i=1}^{N} a_i \tau_i$ where $a_i$ represents the pre-exponential factor and $\tau_i$ the associated fluorescence lifetime.

**Table S2.** Pre-exponential factors, lifetimes and fitting errors associated with vesicles composed of 0.5 % and 0.05 % DiD as a function of excitation power. Experimental conditions: Tris-HCl buffer, pH 7.9, 21°C.

| | **0.04 W/cm²** | | **0.4 W/cm²** | | **0.8 W/cm²** | |
|---|---|---|---|---|---|---|
| [DiD] (molar percent) | **0.5 %** | **0.05 %** | **0.5 %** | **0.05 %** | **0.5 %** | **0.05 %** |
| **$a_1$** | 0.13 ± 0.01 | 0.20 ± 0.01 | 0.09 ± 0.01 | 0.13 ± 0.01 | 0.34 ± 0.01 | 0.38 ± 0.02 |
| **$\tau_1$ / s** | 24.2 ± 0.3 | 18.7 ± 0.3 | 1.8 ± 0.1 | 3.6 ± 0.1 | 3.1 ± 0.1 | 14.2 ± 0.7 |
| **$a_2$** | 0.69 ± 0.01 | 1.04 ± 0.01 | 0.50 ± 0.01 | 0.68 ± 0.01 | 0.61 ± 0.01 | -- |
| **$\tau_2$ / s** | 155.6 ± 7.3 | 447 ± 9.8 | 8.4 ± 0.1 | 35.7 ± 3.3 | 13.8 ± 0.1 | -- |
| **$a_3$** | -- | -- | 0.43 ± 0.01 | -- | -- | -- |
| **$\tau_3$ / s** | -- | -- | 34.4 ± 5.7 | -- | -- | -- |
| **$\tau_{av}$*/ s** | ***151.9 ± 7.1*** | ***438.6 ± 9.7*** | ***28.4 ± 4.4*** | ***35.1 ± 3.2*** | ***12.6 ± 0.1*** | ***14.2 ± 0.7*** |

* Amplitude-weighted average lifetimes were calculated according to the equation $\tau_{av} = \sum_{i=1}^{N} a_i \tau_i^2 / \sum_{i=1}^{N} a_i \tau_i$ where $a_i$ represents the pre-exponential factor and $\tau_i$ the associated decay lifetime.

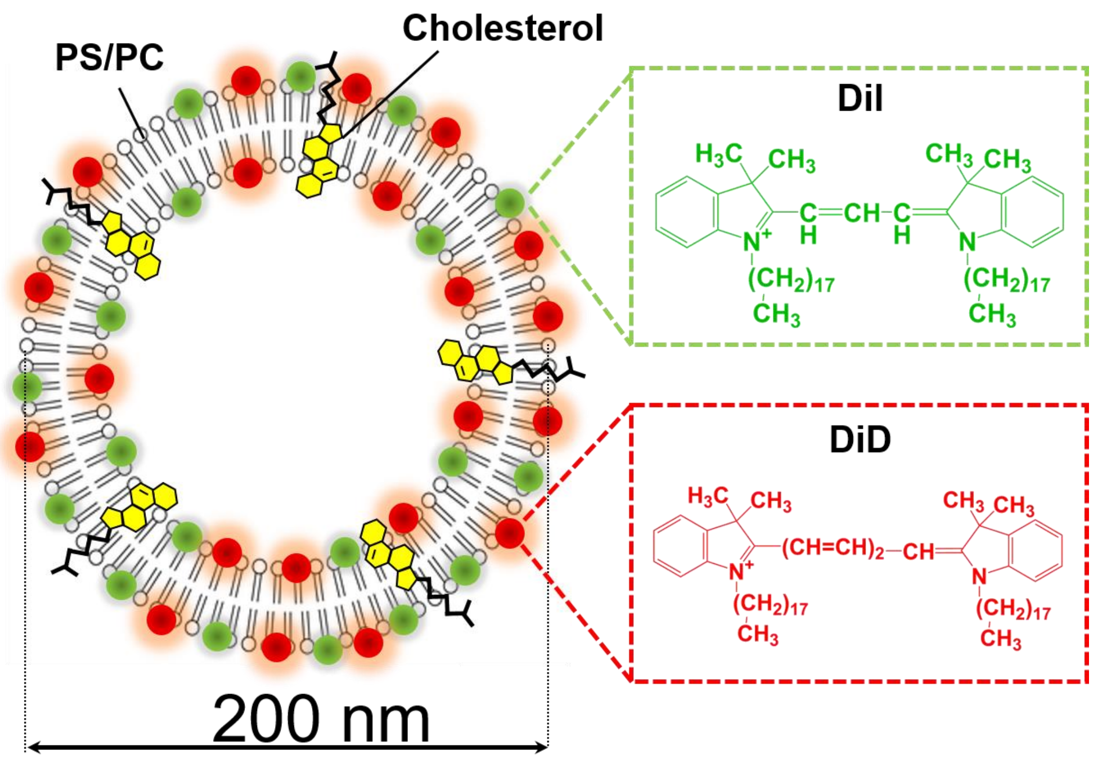


**Figure S1.** Schematic representation of PS/PC vesicles fluorescently-labelled with DiI and DiD. Insets: chemical structures of DiI (green) and DiD (red). DiI, acting as the FRET donor, and DiD, acting as the FRET acceptor are derivatives of the widely used Cy3 and Cy5 dyes (Ro ~ 53 Å) in which the chain between aromatic groups has been extended to enable their incorporation within a lipid bilayer.

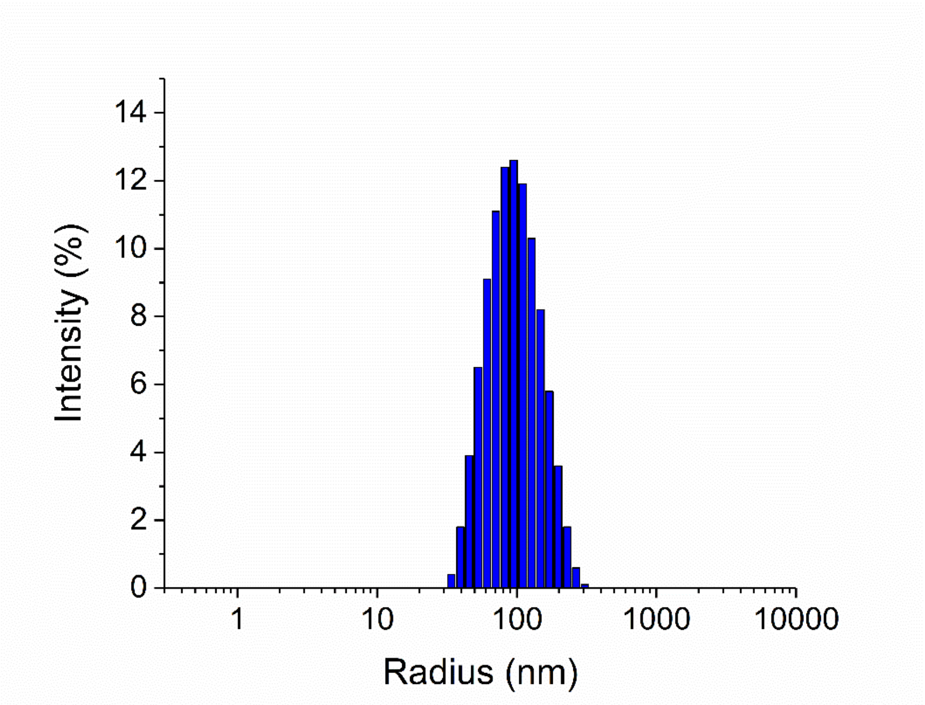


**Figure S2**. Semi-log plot of vesicle radius as a function of scattering intensity, as obtained from dynamic light scattering. Experimental conditions: Tris-HCl buffer, pH 7.9, 21°C.

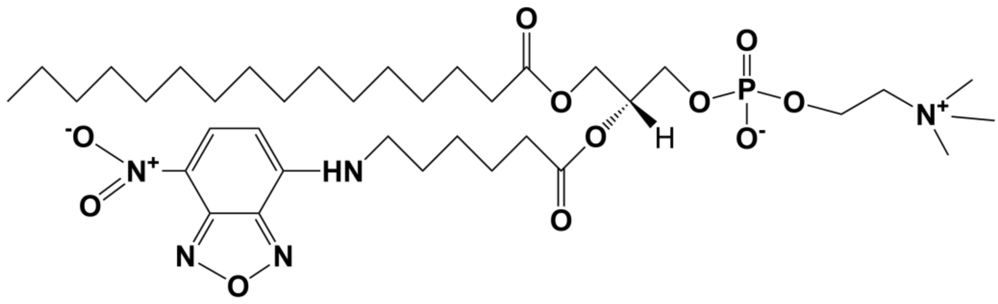


**Figure S3**. Chemical structure of NBD-PC.

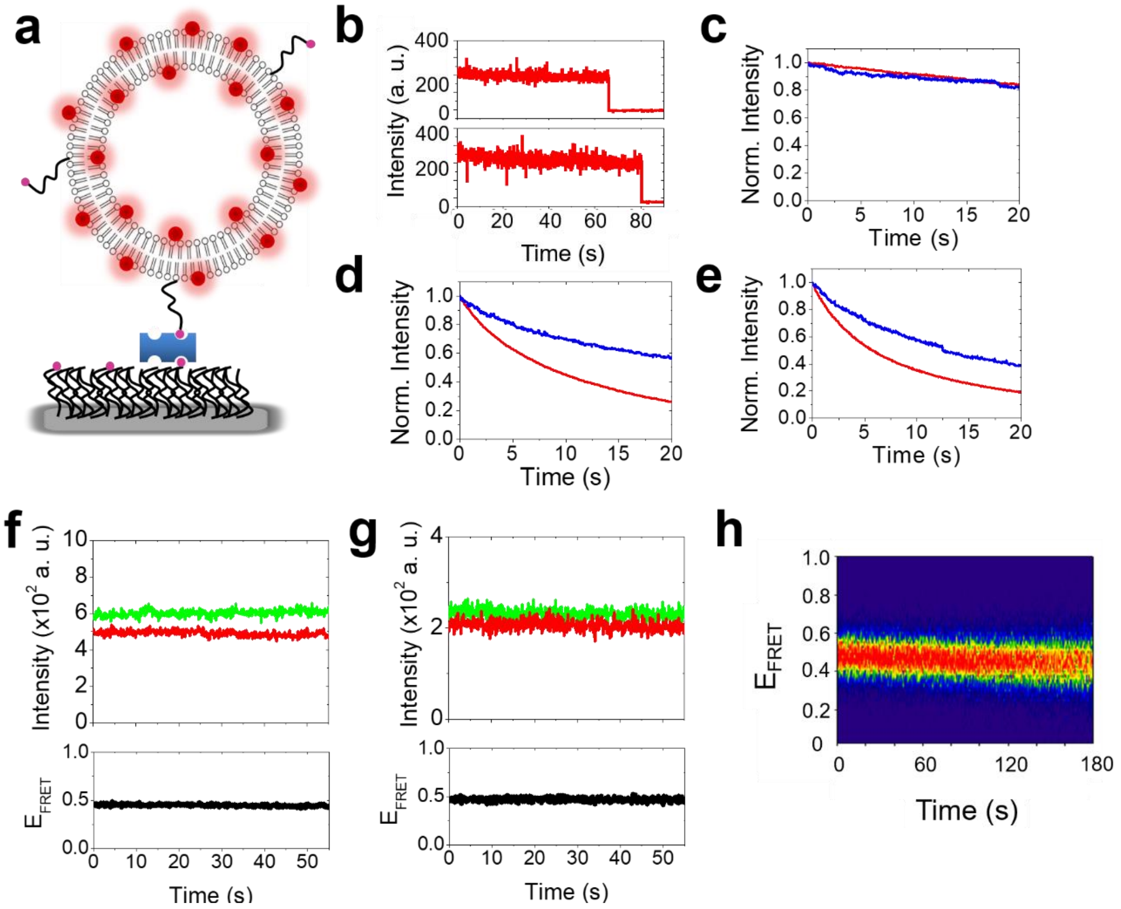


**Figure S4. Optimization of vesicle photostability.** (a) Schematic of the immobilization scheme. (b) Representative stepwise photobleaching trajectories obtained from Cy5-labelled ssDNA under continuous excitation of 0.4 mW/cm$^2$ with $\lambda_{exc}$ = 635 nm. The normalised sum of intensity trajectories from N >1000 individual vesicles labelled with 0.5 % (red) and 0.05 % (blue) DiD are shown for excitation intensities of (c) 0.04 mW/cm$^2$, (d) 0.4 mW/cm$^2$ and (e) 0.8 mW/cm$^2$ ($\lambda_{exc}$ = 635 nm). (f, g) Representative fluorescence trajectories from SUVs composed of 0.125 % DiI (green) and 0.125 % DiD (red) with an excitation intensity < 0.04 mW/cm$^2$. Also shown are the corresponding FRET trajectories ($\lambda_{exc}$ = 532 nm). (h) FRET contour plot extracted from N > 500 vesicles as described with an excitation intensity < 0.04 mW/cm$^2$.

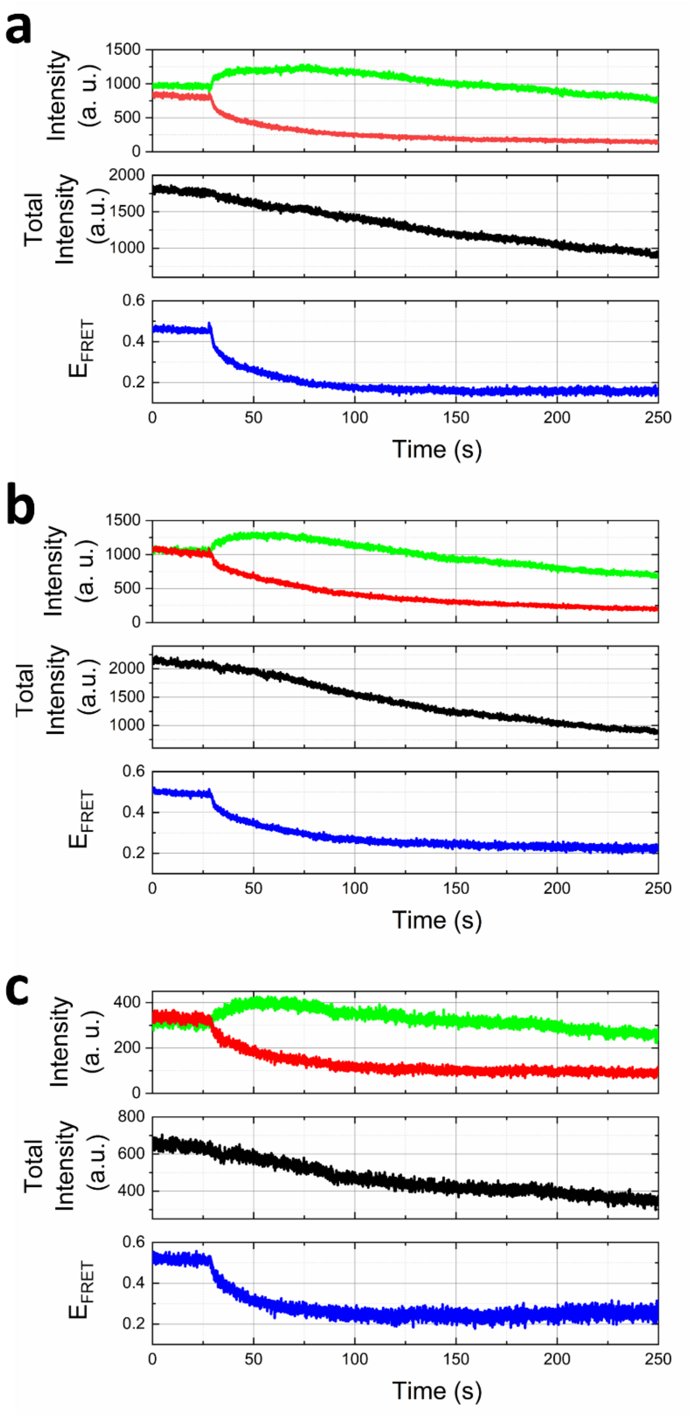


**Figure S5. Real-time visualization of TX-100 induced vesicle solubilization kinetics by svFRET.** (a-c) Representative variation in the fluorescence emission of DiI (green) and DiD (red) (top panel), the sum of their intensities (middle panel) and the corresponding variation in FRET efficiency obtained before (< 25 s) and after (> 25 s) injection of 0.16 mM TX-100 into immobilized vesicles containing 20 % cholesterol.

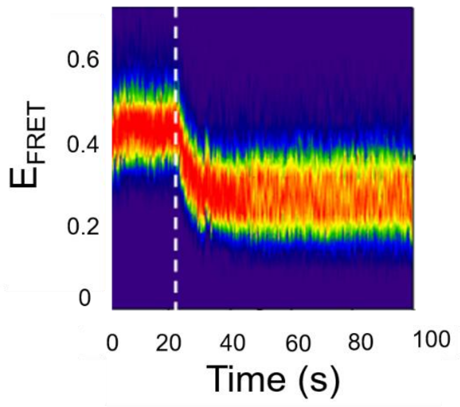


**Figure S6 TX-100 induced vesicle expansion probed by svFRET**. FRET contour plot showing the variation in FRET efficiency as a function of time when 0.16 mM TX-100 is injected (dashed white line) into immobilized DiI/DiD labelled PS/PC vesicles in the absence of cholesterol.
